# Genome Wide Association Scan identifies new variants associated with a cognitive predictor of dyslexia

**DOI:** 10.1101/309336

**Authors:** Alessandro Gialluisi, Till F M Andlauer, Nazanin Mirza-Schreiber, Kristina Moll, Per Hoffmann, Kerstin U Ludwig, Darina Czamara, Clyde Francks, Beate St Pourcain, William Brandler, Ferenc Honbolygó, Dénes Tóth, Valéria Csépe, Guillaume Huguet, Andrew P Morris, Jacqueline Hulslander, Erik G Willcutt, John C DeFries, Richard K Olson, Shelley D Smith, Bruce F Pennington, Anniek Vaessen, Urs Maurer, Heikki Lyytinen, Myriam Peyrard-Janvid, Paavo H T Leppänen, Daniel Brandeis, Milene Bonte, John F Stein, Joel B Talcott, Fabien Fauchereau, Thomas Bourgeron, Anthony P Monaco, Franck Ramus, Karin Landerl, Juha Kere, Thomas S Scerri, Silvia Paracchini, Simon E Fisher, Johannes Schumacher, Markus M Nöthen, Bertram Müller-Myhsok, Gerd Schulte-Körne

## Abstract

Developmental dyslexia (DD) is one of the most prevalent learning disorders among children and is characterized by deficits in different cognitive skills, including reading, spelling, short term memory and others. To help unravel the genetic basis of these skills, we conducted a Genome Wide Association Study (GWAS), including nine cohorts of reading-impaired and typically developing children of European ancestry, recruited across different countries (N=2,562-3,468).

We observed a genome-wide significant effect (p<1×10^−8^) on rapid automatized naming of letters (RANlet) for variants on 18q12.2 within *MIR924HG (micro-RNA 924 host gene*; *p* = 4.73×10^−9^), and a suggestive association on 8q12.3 within *NKAIN3* (encoding a cation transporter; *p* = 2.25 ×10^−8^). RAN represents one of the best universal predictors of reading fluency across orthographies and linkage to RAN has been previously reported within *CELF4* (18q12.2), a gene highly expressed in the fetal brain which is co-expressed with *NKAIN3* and predicted to be a target of *MIR924*. These findings suggest new candidate DD susceptibility genes and provide insights into the genetics and neurobiology of dyslexia.

## Background

Developmental dyslexia (DD) is a neurodevelopmental disorder affecting the ability of learning to read, in spite of adequate intelligence, educational opportunities and in the absence of overt neurological and sensorial deficits^1^. It shows a prevalence of 5–12% among school-aged children, implying life-long learning difficulties for most of the affected individuals^1^. Dyslexic individuals usually show problems in accurate and fluent reading and spelling, and in reading comprehension^2^. These problems are often caused by deficits in underlying cognitive skills, such as the ability to recognize and manipulate the phonemic constituents of speech (also known as phoneme awareness), the ability to store such phonemes while reading (also known as phonological short term memory), or the ability to fast map known visual symbols onto spoken word representations (known as naming speed)^3^. All these cognitive abilities show moderate to high heritability (40–80%)^4–6^ and significant genetic correlations with DD^4^. Hence, they represent cognitive indicators of dyslexia risk that are optimally suited for investigating the genetic mechanisms at its basis.

In the last two decades, several studies investigating both DD and the underlying cognitive skills have been carried out to better understand the genetic and neurobiological basis of reading. On the one hand, linkage and targeted association analyses have suggested several candidate DD susceptibility genes, the most robust of which include *DYX1C1* (15q21), *DCDC2* and *KIAA0319* (6p22.3), *GCFC2* and *MRPL19* (2p12), and *ROBO1* (3p12.3-p12.3) (reviewed in ^1,7,8^). On the other hand, most of the genome-wide association studies (GWAS) published so far have identified mainly suggestive associations with DD and related cognitive traits (*p* < 10^−5^)^9–13^, with only one recent study reporting a genome-wide significant association (*p* < 5×10^−8^)^14^. The first GWAS for reading ability reported used DNA pooling of low vs high reading ability groups in ~1,500 7-year-old children which were genotyped with a low-density Single Nucleotide Polymorphism (SNP) microarray (~ 107,000 SNPs)^13^. The SNPs showing the largest allele frequency differences between low and high ability groups were tested in an additional follow-up cohort of 4,258 children, finally identifying ten SNPs showing nominally significant associations with continuous variation in reading ability^13^. A later genome-wide linkage and association scan on ~133,000 SNPs, in a sample of 718 subjects from 101 dyslexia-affected families, identified an association with dyslexia status at rs9313548, near *FGF18* (5q35.1)^12^. More recently, three GWAS studies with different designs were carried out with the aim of identifying shared genetic contributions to reading and language abilities. Luciano et al.^11^ performed a GWAS on quantitative reading- and language-related traits in two population-based cohorts (N~6,500), analysing word reading, nonword repetition, and a composite score of reading and spelling abilities. They reported a suggestive association of rs2192161 (*ABCC13;* 21q11.2) with nonword repetition and of rs4807927 (*DAZAP1*, 19p13.3) with both the word reading and the reading-spelling score. A case-control GWAS comparing dyslexic (N=353), language impaired (LI) (N=163), and comorbid cases (N=174) to a population-based control dataset (N=4,117) identified nominally significant associations with comorbid DD-LI cases at rs12636438 and rs1679255, mapping to *ZNF385D* (3p24.3)^9^. Another GWAS analysed the first principal component from various reading- and language-related traits (both with and without IQ adjustment) in three datasets comprising children with reading or language problems and their siblings (N=1,862), and reported suggestive associations at rs59197085, upstream of *CCDC136/FLNC* (7q32.1), and at rs5995177, within *RBFOX2* (22q12.3)^10^. More recently, Truong and colleagues^14^ reported a genome-wide significant multivariate association of rs1555839 (10q23.31) with two skills predicting DD risk, namely rapid automatized naming (RAN) and rapid alternating stimulus (RAS), in a multisite case-control study of DD made up of individuals of non-European ancestry (N=1,263). This SNP, located upstream of the pseudogene *RPL7P34*, was also associated with measures of word reading and showed a nominally significant multivariate association (*p* < 0.05) with RAN traits in an independent cohort from Colorado.

Although many of the genes suggested by these GWAS studies showed interesting potential biological links to DD and underlying skills, most of these associations did not reach genome-wide significance and were not replicated in independent datasets^15^. This might have different reasons, including the low statistical power of these studies implied by the relatively small sample sizes, and the heterogeneity of recruitment criteria and phenotypic assessment of the cohorts involved. In addition, the candidate susceptibility genes identified and replicated so far explain only a minor part of the genetic variance underlying dyslexia and the related cognitive traits, and a big proportion of this heritability remains unexplained.

To help unravel the genetic basis of DD and related cognitive skills, we conducted a large international collaborative GWAS. We analysed cognitive traits such as word reading, spelling, decoding skills, phoneme awareness, verbal short term memory and naming speed, in nine cohorts of reading impaired and typically developing participants of European ancestry (maximum N=3,468). We observed a genome-wide significant association at 18q12.2 and an association approaching genome-wide significance at 8q12.3, both with rapid automatized naming (RAN, N=2,563), which allowed us to identify two novel candidate susceptibility genes potentially affecting this ability.

## Subjects and Methods

### Datasets

Table 1 reports the main details on the datasets involved in this study and on the recruitment criteria.

**Table 1.**
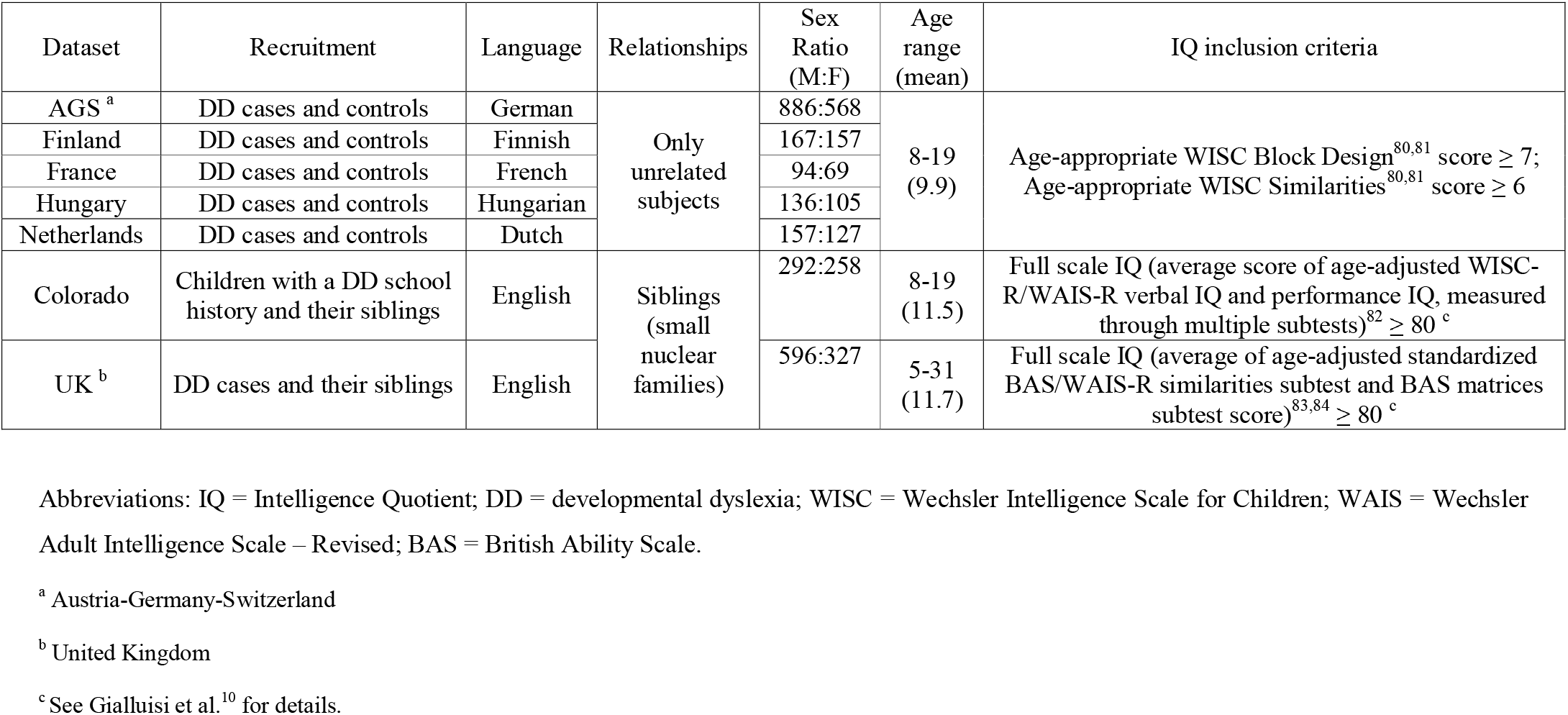
Main characteristics and recruitment criteria of the datasets involved in the present study.

Unrelated DD cases and controls were recruited across seven different European countries, namely Austria (N=374), Germany (N=1,061), Finland (N=336), France (N=165), Hungary (N=243), The Netherlands (N=311), and Switzerland (N=67). In addition, we included two family-based datasets in the study. One of these, from Colorado, United States (US), contained children showing a school history of reading difficulties as well as their siblings (N=585; 266 independent nuclear families)^10,16^. The other one, from the United Kingdom (UK), consisted of subjects with a formal diagnosis of dyslexia and their unaffected siblings (N=983; 608 independent nuclear families)^10,17^. Although the family-based datasets have been previously investigated in GWAS studies^10,17,18^ and the European datasets have been analysed in a candidate (SNP) association study^19^, such datasets were never analysed jointly in a GWAS. In the present study, samples from Austria, Germany, and Switzerland were merged into a single dataset (hereafter called AGS), since they shared language, genetic ancestry, phenotypic measures and selection criteria^19–21^.

### Phenotypic measures

We focused on the core phenotypes of dyslexia, namely word reading (WRead), nonword reading (NWRead), and word spelling (WSpell), and on five cognitive measures underlying reading ability and dyslexia, namely phoneme awareness (PA), digit span (DigSpan, a measure of verbal short-term memory), and rapid automatized naming of letters (RANlet), digits (RANdig), and pictures (RANpic). These skills showed moderate to high cross-trait correlations (see Table S1 in *Supplementary Methods*). A brief explanation of these measures is reported in Table 2, while details on statistical elaboration are reported in *Supplementary Methods* and elsewhere ^10,20,21^. Briefly, raw scores from psychometric tests were grade-normed (age-adjusted in Colorado) and then z-standardized to reduce skewness, with the exception of the DigSpan score, which was only z-normalized in all datasets since it was already standardized and normally distributed^20^. No phenotypic outliers were detected in any of the datasets analysed (see *Supplementary Methods* for details).

**Table 2.** Cognitive traits analysed in the present study. More detailed information on these phenotypic measures, including psychometric tests used and statistical elaboration, is reported in the *Supplementary Methods*.

| Trait | Definition | Task |
| --- | --- | --- |
| Wread | Reading single real words of varied difficulty | Timed word reading in AGS, Finland, France, Hungary and the Netherlands;<br>Untimed word reading in UK;<br>Composite score of timed word reading and reading accuracy in Colorado |
| Wspell | Spelling single real words after dictation | Spelling accuracy |
| NWRead | Reading aloud nonsense words of varied difficulty | Timed nonword reading in AGS, Finland, France, Hungary and the Netherlands;<br>Untimed nonword reading in UK and Colorado |
| PA | Deletion, substitution or swapping of specific phonemes in one or multiple words | Phoneme deletion in AGS, Finland, France, Hungary and the Netherlands;<br>Phoneme deletion/substitution and spoonerism in UK;<br>Composite of phoneme deletion and phoneme segmentation and transposition tasks in Colorado |
| DigSpan | Reciting a sequence of digits presented by recalling them in the same (forward) and/or reverse (backward) order | WISC (Wechsler Intelligence Scale for Children) forward and backward digit span task |
| RANdig | Naming as quickly and as accurately as possible a matrix of digits visually presented | Naming speed task (number of digits correctly named per minute) |
| RANlet | Naming as quickly and as accurately as possible a matrix of letters visually presented | Naming speed task (number of letters correctly named per minute) |
| RANpic | Naming as quickly and as accurately as possible a matrix of objects visually presented | Naming speed task (number of objects/pictures correctly named per minute) |

### Genotype quality control (QC) and imputation

Individuals were genotyped using Illumina HumanHap 300k, 550k, 660k, HumanOmniExpress, and HumanCoreExome BeadChips (see Table S2 for details). Genotype QC was carried out in PLINK v1.90b3s (https://www.cog-genomics.org/plink2)^22^ and QCTOOL v1.4 (http://www.well.ox.ac.uk/~gav/qctool/), as described in *Supplementary Methods* and elsewhere^23^. Within each dataset, SNPs were filtered out if they showed a variant call rate < 98 %; a minor allele frequency (MAF) < 5%, or a Hardy Weinberg Equilibrium (HWE) test *p*-value < 10^−6^. Moreover, samples showing a genotyping rate < 98 %, cryptic relatedness (in datasets of unrelated subjects), identity-by-descent (IBD) not corresponding to the available pedigree information (in sibling-based datasets), and mismatches between genetic and pedigree-based sex were discarded. Furthermore, genetic ancestry outliers-detected in a multidimensional scaling (MDS) analysis of pairwise genetic distance- and samples showing significant deviations in genome-wide heterozygosity were also filtered out (see Table S3).

For imputation, autosomal variants were aligned to the 1000 Genomes phase I v3 reference panel (ALL populations, June 2014 release)^24^ and pre-phased using SHAPEIT v2 (r837)^25^. Imputation was performed using IMPUTE2 v2.3.2^26^ in 5 Mb chunks with 500 kb buffers, filtering out variants that were monomorphic in the 1000 Genomes EUR (European) samples. Chunks with < 51 genotyped variants or concordance rates < 92 % were fused with neighboring chunks and re-imputed. Finally, imputed variants (genotype probabilities) were filtered for IMPUTE2 INFO metric ≥ 0.8, MAF < 5% and HWE test *p*-values < 10^−6^, using QCTOOL v1.4. We checked again for the absence of genetic ancestry and genome-wide heterozygosity outliers after imputation, which revealed substantial concordance with preimputation QC. Further details on the filters used in genotype QC are reported in Table S3, while summary statistics are reported for each dataset in Table S2.

### Genetic association testing and meta-analysis

After genotype QC and imputation, autosomal genotype probabilities were tested for association with the continuous traits available within each dataset. In the datasets containing only unrelated subjects – namely AGS, Finland, France, Hungary, and The Netherlands – association with genotype dosage was tested through linear regression in PLINK v1.9, using the first ten genetic ancestry (MDS) components as covariates. In the sibling-based datasets (Colorado and UK), a generalized linear mixed-effects model association test was carried out through FastLMM v2.07^27^, using a genetic relationship matrix (GRM) of samples as a random effect while disabling normalization to unit variance for tested SNPs.

Following separate GWAS analyses for each dataset, variant associations with each of the eight univariate traits available were combined using a fixed-effects model based on inverse-variance-weighted effect size in METASOFT v2.0.1^28^. Following the software guidelines, pooled analysis was conducted in two steps: a first run was carried out to compute genomic inflation factors, which were then used to correct meta-analysis statistics in a second run. The numbers of subjects involved in our pooled analysis were 3,468 for WRead, 3,399 for WSpell, 3,409 for NWRead, 3,093 for PA, 2,591 for DigSpan, 2,563 for RANlet and RANdig, and 2,562 for RANpic (see Table S4 for detailed sample size by dataset). RAN measures and DigSpan were not available in the UK dataset, which was therefore not included in the pooled analyses of those traits. The numbers of variants analysed in two or more datasets were 6,952,813 for RANlet, RANdig, RANpic, and DigSpan and 6,969,139 for WRead, WSpell, NWRead, and PA. The common genome-wide significance threshold α = 5×10^−8^ was corrected for multiple testing of five independent latent variables, as computed through MatSpD (http://gump.qimr.edu.au/general/daleN/matSpD/)29 on the correlation matrix of the eight univariate traits analysed (Table S1). This adjustment resulted in a final Bonferroni-corrected significance level α = 1×10^−8^.

### Further analyses of top association signals

The analyses explained in this section were only conducted on datasets with RANlet measures available (see Table S4) and required the preliminary adjustment of phenotypic traits for genetic population structure in each dataset. This was carried out differently in the datasets including only unrelated subjects (AGS, Finland, France, Hungary, and The Netherlands) and in the sibling-based dataset (Colorado). In the former group, we regressed the phenotypic traits against the first ten MDS components (previously used as covariates in the GWAS). In the latter case, we adjusted the traits for a GRM through the *polygenic()* function of the GenABEL package (http://www.genabel.org/)^30^.

### Permutation-based correlation test and effect size estimation

To assess the robustness of the most significant associations detected (with RANlet), we carried out a permutation-based test on the top-associated SNPs at 18q12.2 (rs17663182) and 8q12.3 (rs16928927) in R v3.2.3 (http://www.R-project.org/)31. Briefly, we first computed allelic dosages from genotype probabilities for the SNPs of interest within each dataset, and adjusted the RANlet score for genetic population structure in each dataset (as explained above). Subsequently, we computed Pearson correlation through the *cor()* function of the WGCNA v1.51 package^32^. After the calculation of the Pearson correlation coefficient r, we permuted both phenotypic residuals and dosages 10,000 times, computing similar correlation coefficients for each of the resulting 10,000 × 10,000 = 100 million random combinations. Finally, we derived an empirical *p*-value from the distribution of these 100 million random correlations (defined as the frequency of random correlations which were at least as high as our original correlation coefficient r).

To estimate the fraction of RANlet phenotypic variance explained by rs17663182 (18q12.2) and rs16928927 (8q12.3) within each dataset, we used R to compute linear regression *R*^2^ of the phenotypic trait adjusted for genetic population structure vs dosage values of the top-associated variants.

### Test of pleiotropy

We tested the top association signals for pleiotropic effects on traits other than RANlet analysed in this study, namely WRead, WSpell, NWRead, PA, DigSpan, RANdig and RANpic. To this end, we first regressed these traits, which had previously been adjusted for genetic population structure, against the RANlet score in R, separately for each dataset. Then we tested the residuals of these traits for association with rs17663182 and rs16928927 dosages in PLINK. Finally, we combined the results of the association tests in different datasets through an inverse-variance fixed-effect pooled analysis in METAL v25-03-2011 (http://www.sph.umich.edu/csg/abecasis/Metal/index.html)33, which allowed us to directly detect concordance of allelic trends across datasets for all the SNPs tested.

### Test for independent genetic effects in 18q12.2 and 8q12.3

We tested for the presence of genetic effects independent from the local top hits in 18q12.2 and 8q12.3 (see above). For each of these two SNPs, we first regressed RANlet scores adjusted for population structure against the allelic dosage values and extracted the phenotypic residuals in each dataset. Then we used PLINK v1.9 to test these residuals for association with all the SNPs positioned up to 50 kb from the most significant variant in each region of interest, namely 275 variants on 8q12.3 and 236 variants on 18q12.2. Then we combined the association statistics that were produced for each dataset using METAL (as described above).

### SNP× SNP interaction analysis

To investigate potential epistatic effects of rs17663182 and rs16928927 on RANlet, we carried out a two-SNP interaction analysis in R. Since rs16928927 was not available in the Finnish dataset, this analysis was conducted only in the AGS, France, Hungary, Netherlands, and Colorado datasets. The analysis consisted of two steps: first, we regressed RANlet scores adjusted for genetic population structure against the allelic dosages of the SNPs rs17663182 and rs16928927. Then we regressed the RANlet residual scores against a single interaction term of the two SNPs and computed the fraction of phenotypic variance (*R*^2^) explained by this term.

### Imaging genetics follow-up

To further investigate the potential neurobiological implications of the top association signals detected at rs17663182 (18q12.2) and rs16928927 (8q12.2), we assessed genetic effects of these SNPs on different subcortical volumes, including Nucleus Accumbens, Amygdala, Caudate Nucleus, Hippocampus, Pallidum, Putamen and Thalamus. These neuroimaging traits had been tested for association in a large GWAS involving 30,717 subjects of European ancestry^34^. Our choice of investigating subcortical brain volumes was determined by two factors, namely i) the increasing evidence implicating subcortical structures in reading and language abilities (as reviewed in ^1,35,36^), and ii) the large sample size of the imaging genetics GWAS, which maximized the power to detect significant genetic effects.

For this analysis, we computed a Bonferroni-corrected significance threshold α = 7.1×10^−4^, taking into account two SNPs, five independent latent traits tested in our study (computed in MatSpD, see above), and the seven neuroimaging subcortical regions analysed by Hibar and colleagues^34^.

### Assessment of genes and SNPs previously associated with DD and related cognitive traits

We investigated single variant associations for candidate SNPs and genes previously implicated in DD and related cognitive traits.

First, we assessed all the variants mapping to nine candidate genes (up to 10 kb from the 5’-or 3-UTR): *DYX1C1, DCDC2, KIAA0319, C2ORF3, MRPL19, ROBO1, GRIN2B, FOXP2* and *CNTNAP2*. For these genes, association with DD and related cognitive traits was previously reported in at least two independent studies (as reviewed in ^1^). Of note, most of the candidate variants identified in these genes have been already tested in studies showing a variable degree of overlap with our cohorts (reviewed in ^1,7,8^) hence they cannot be formally replicated within the scope of the current study. For this reason, we focused on six candidate SNPs among these variants, for which a statistically significant association (p < 0.05 after correction for multiple testing) has been reported in the past in datasets other than ours, but was never formally replicated. These SNPs included rs6803202, rs4535189, rs331142 and rs12495133 in ROBO1^37,38^, rs7782412 in *FOXP2*^39^ and rs5796555 in *GRIN2B*^40^.

We next tested all the variants showing the strongest associations with DD and related cognitive traits in previous GWAS^9–14^. These included all those variants reported to be associated in previous GWAS papers, including genome-wide significant associations (*p* < 5×10^−8^), suggestive associations (p < 1×10^−5^), or variants reported as the most significant associations (top 10 or top 100 list, depending on the paper; see *Results* section for a complete list). Again, some of these variants were identified by studies partially overlapping with our datasets^10^, while for other SNPs tested the statistics from the original papers were not fully available or not always directly comparable, due to either different design of the study or to different traits analysed^9–14^. Therefore, a direct comparison was possible only for few variants (see relevant *Results* section).

### Gene- and pathway-based enrichment tests

Gene-based association analyses for the phenotypic traits analysed were performed using MAGMA v1.06 (http://ctg.cncr.nl/software/magma)41. First, genetic variants were assigned to protein-coding genes based on their position according to the NCBI 37.3 (hg19) build, extending gene boundaries by 10kb from the 3’- and 5’-UTR. A total of 18,033 genes (out of 19,427 genes available) included at least one variant that passed internal QC, and were thus tested in gene-based enrichment analysis. Gene-based statistics were computed using the single-variant association statistics calculated in the GWAS of each phenotype, using default settings. To account for linkage disequilibrium (LD) among the variants tested, we used a combined genetic data set of all the datasets pooled together. Given the number of genes (18,033) and of independent latent traits (5) tested, the Bonferroni corrected genome-wide significance threshold for this analysis was set to α = 0.05 / (18,033 × 5) = 5.5×10^−7^.

Using the results of the gene-based association analysis, we carried out a pathway-based enrichment test for each trait analysed in the study, through a competitive gene-set analysis in MAGMA v1.06. We tested for enrichment 1,329 canonical pathways (i.e. classical representations of biological processes compiled by domain experts) from the Molecular Signatures Database website (MSigDB v5.2; http://software.broadinstitute.org/gsea/msigdb; *collection C2, subcollection CP)*. To correct enrichment statistics for testing of multiple pathways, we used an adaptive permutation procedure with default settings (up to a maximum of 10,000 permutations). Hence, for gene-set analysis we corrected the significance threshold only for the number of independent latent traits tested (α = 0.05 / 5 = 0.01).

### Polygenic Risk Score analysis

To assess the genetic overlap of common variants between the dyslexia-related skills tested here and other correlated phenotypes, we carried out a polygenic risk score (PRS) analysis using PRSice v1.25^42^. We used the eight traits analysed in our GWAS as target traits and selected twelve different training traits from previous GWAS studies, including seven subcortical volumes used for the imaging genetics follow-up^34^, an educational attainment trait (expressed in years of education completed, EDUyears; N~293,000)^43^ and four neuropsychiatric disorders. These included Attention Deficit Hyperactivity Disorder (ADHD; N~55,000)^44^; Autism Spectrum Disorder (ASD; N~16,000)^45^; Major Depressive Disorder (MDD; N~19,000)^46^; and Schizophrenia (SCZ; N~150,000)^47^. These neuropsychiatric conditions were selected in light of their comorbidity with dyslexia reported by previous literature^3,48-50^.

We performed an analysis of summary statistics using only SNPs with association *p*-values ≤ 0.05, and in linkage equilibrium (r^2^ < 0.05) with the local top hit within a 300 kb window, in each training GWAS. Only SNPs which had been tested both in the training and in the target GWAS were tested. The number of SNPs meeting these criteria ranged from 11,017 for the comparison of MDD vs DigSpan and RAN traits, to 25,409 for SCZ vs WRead, WSpell, NWRead and PA. To verify the robustness of our results, we repeated the analysis at increasing association significance (*P_T_*) thresholds in the training GWAS (with *P_T_* = 0.001, 0.05, 0.1, 0.2, 0.3, 0.4, 0.5).

To obtain a statistic on the direction of genetic correlations, we selected variants with association *p*-values ≤ 0.05 in each training GWAS and computed Pearson’s correlation of effect sizes (hereafter called r_β_) with each of the target GWAS analysed. The significance threshold for these analyses was corrected for multiple testing of five independent target GWAS (i.e. the number of independent latent traits computed through MatSpD) and twelve different training GWAS (α = 0.05 / (5 × 12) = 8.3×10^−4^.

## Results

For each analysis presented below, we report the empirical p-values, along with significance thresholds adequately corrected for multiple testing (see *Subjects and Methods* section).

### Single variant genome-wide associations

Among the eight traits analysed in the present GWAS, only RANlet showed genome-wide significant associations withstanding correction for multiple testing (*p* < 1×10^−8^), mapped to chromosome 18q12.2. The most significant association was observed for rs17663182 (G/T; MAF = 7.7%; *p*-value = 4.73×10^−9^, major allele (G) β (SE) = 0.35 (0.06)). All the SNPs significantly associated on 18q12 were located within the non-coding gene *MIR924HG (micro-RNA 924 host gene*, also known as *LINC00669*; see Figure 1a) and were in high LD with each other (*r*^2^ > 0.9). An additional, independent association approaching genome-wide significance was observed with RANlet at rs16928927 (C/T; MAF = 6.5%; *p*-value = 2.25×10^−8^, major allele (C) β (SE) = −0.4 (0.07)) on 8q12.3. This SNP was located within the first intron of *NKAIN3* (*Na+/K+ transporting ATPase interacting 3*; see Figure 1b). Further details on these associations are reported in Figure 2 and Table 3, while more detailed results of the GWAS analyses for each trait are reported in Supplementary Figures S1a-p and Tables S5a-h.

**Figure 1.**
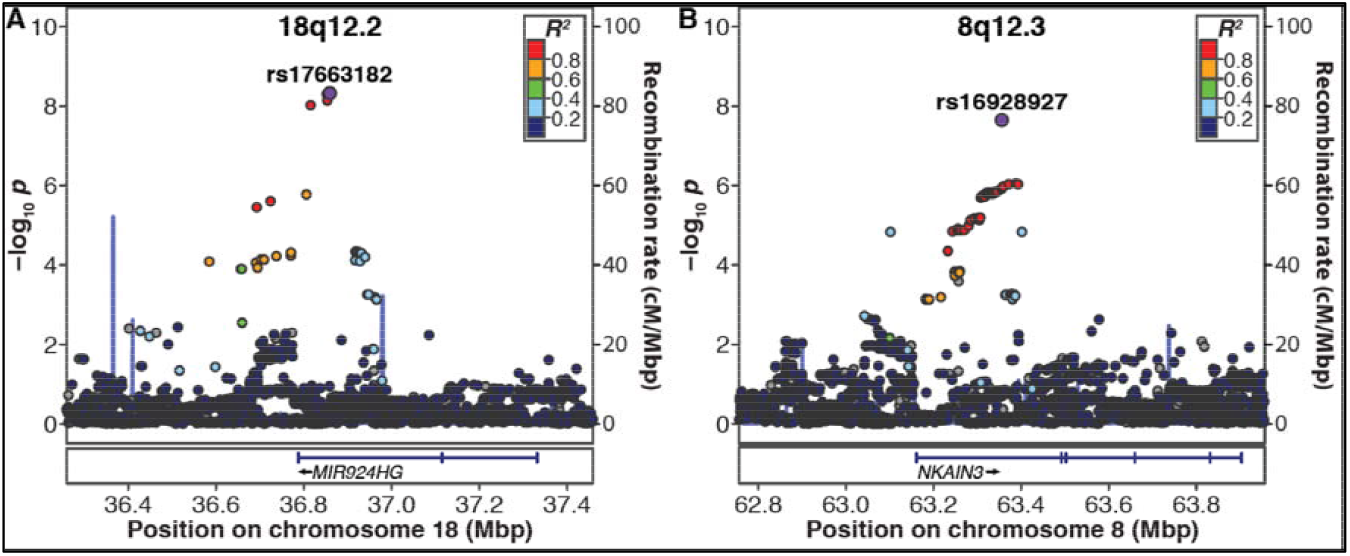
Regional association plots of **a)** 18q12.2 and **b)** 8q12.3 with the RANlet trait. The most significantly associated variants are highlighted in violet. Plots were made using LocusZoom (http://www.locuszoom.org/).

**Figure 2.**
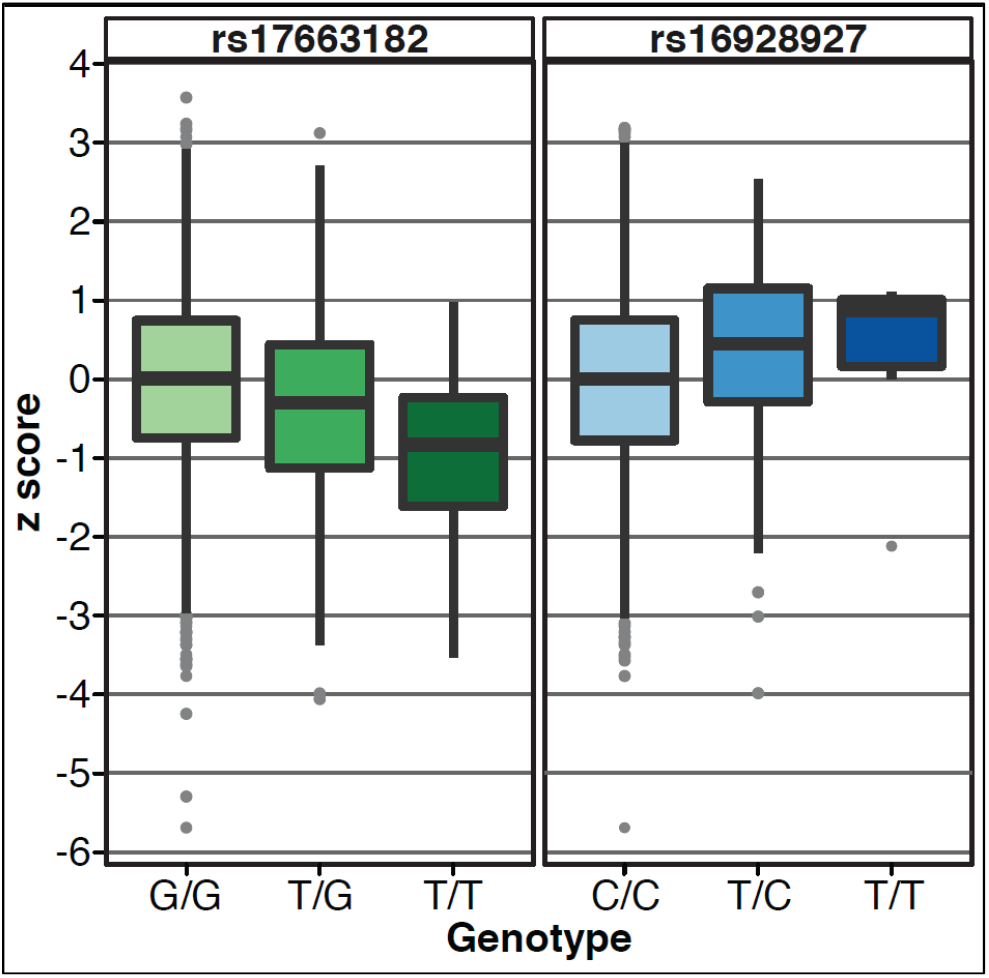
Boxplots of the RANlet trait as a function of genotype of the lead variants rs17663182 (left side, major allele G) and rs16928927 (right side, major allele C). To generate these plots, all datasets were pooled together. RANlet Z-scores plotted here are residualized against the first 10 MDS covariates in all datasets except for Colorado, where we adjusted the phenotypic measure for pairwise genetic relatedness in GenABEL (see *Subjects and Methods* section).

**Table 3.** Most significant single variant associations (p < 1 × 10^−7^) detected in the eight GWAS analyses of the present study.

| SNP | A1 | A2 | A1 frequency <sup>a</sup> | *p*-value | $\beta$ <sup>b</sup> | $\beta$ SE | I<sup>2</sup> <sup>c</sup> | Location (chr:bp) | LD relative to local top hit (*r*<sup>2</sup>) | Gene symbol | Position relative to gene | Distance from gene (bp) | Trait |
| --- | --- | --- | --- | --- | --- | --- | --- | --- | --- | --- | --- | --- | --- |
| rs17663182 | G | T | 0.92 | 4.73 x 10<sup>-9</sup> | 0.353 | 0.060 | 0 | 18:36859202 | - | *LINC00669* | within | - | RANlet |
| rs17605546 | G | A | 0.92 | 4.92 x 10<sup>-9</sup> | 0.352 | 0.060 | 0 | 18:36852398 | 0.98 | *LINC00669* | within | - | RANlet |
| rs74500110 | C | T | 0.92 | 7.14 x 10<sup>-9</sup> | 0.343 | 0.059 | 0 | 18:36853535 | 0.94 | *LINC00669* | within | - | RANlet |
| rs34822091 | G | A | 0.92 | 9.44 x 10<sup>-9</sup> | 0.347 | 0.060 | 0 | 18:36815582 | 0.94 | *LINC00669* | within | - | RANlet |
| rs16928927 | C | T | 0.94 | 2.25 x 10<sup>-8</sup> | -0.403 | 0.072 | 0 | 8:63356625 | - | *NKAIN3* | within | - | RANlet |
| rs1541518 | G | T | 0.71 | 6.42 x 10<sup>-8</sup> | -0.177 | 0.033 | 0 | 7:31148279 | - | *ADCYAP1R1* | downstream | 1956 | NWRead |

<sup>a</sup> Average allele frequency computed over all the datasets analysed.
<sup>b</sup> $\beta$ values are relative to A1.
<sup>c</sup> I-squared test for heterogeneity of genetic effect across datasets (the closer to “0”, the more homogenous is the genetic effect).

### Characterization of top association signals

We examined the top local association signals on 18q12.2 (rs17663182) and 8q12.3 (rs16928927) in detail. Although neither of the two top SNPs was genotyped, imputation quality was high in all datasets (IMPUTE2 INFO metric 0.89-0.94 for rs17663182 and ~0.99 for rs16928927).

Association test statistics showed consistent allelic trends across all datasets for both SNPs (Figure 3a, b and Table S6a, b). Furthermore, the associations were confirmed by an independent permutation-based correlation test between allelic dosages and RANlet scores. Indeed, Pearson correlation *p*-values for both SNPs were very similar to the linear regression *p*-values (see Table S6a, b). The proportion of RANlet variance explained by the SNPs ranged from 0.03% in the Dutch dataset to 1.8% in the AGS dataset for rs17663182 and from 0.067% in AGS to 2.96% in Hungary for rs16928927 (Table S6a, b).

**Figure 3.**
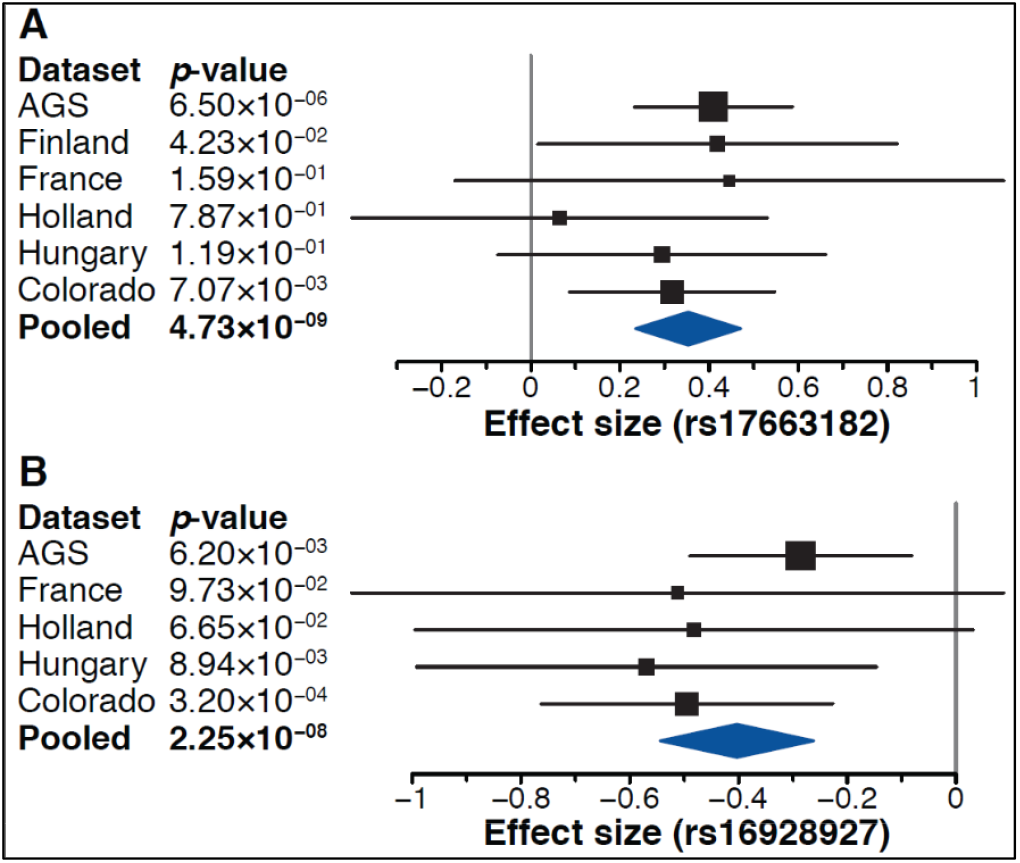
Forest plots of association signals with RANlet for **a)** rs17663182 (18q12.2) and **b)** rs16928927 (8q12.3). Effect sizes (β) refer to major alleles **a)** G and **b)** C, respectively.

Since both our lead SNPs showed evidence of an association with many of the traits analysed in the present study (see Figure 4a, b), we carried out a pleiotropy test: we first regressed the phenotypic traits other than RANlet against this score and then tested the residuals of this model for an association with either rs17663182 or rs16928927 dosages in each dataset separately, followed by fixed-effects pooled analysis. Neither of the two SNPs showed significant effects on any trait other than RANlet (see Tables S6c, d). Similarly, we tested for the presence of independent genetic effects at 18q12.2 and 8q12.3, in a 100 kb window surrounding the two most strongly associated variants. Pooled analysis of association tests with RANlet residual scores extracted from regression against rs17663182 and rs16928927 dosages revealed no independent associations surviving correction for multiple testing (see Tables S6e, f). In addition, we conducted a SNP×SNP interaction analysis on rs17663182 and rs16928927, which revealed no significant epistatic effects of these two SNPs on RANlet: regression *R*^2^ values for the interaction term ranged from 0.6% (*p* = 0.7) in the Dutch dataset to 0.0006% (*p* ~ 1) in the AGS dataset (Table S6g).

**Figure 4.**
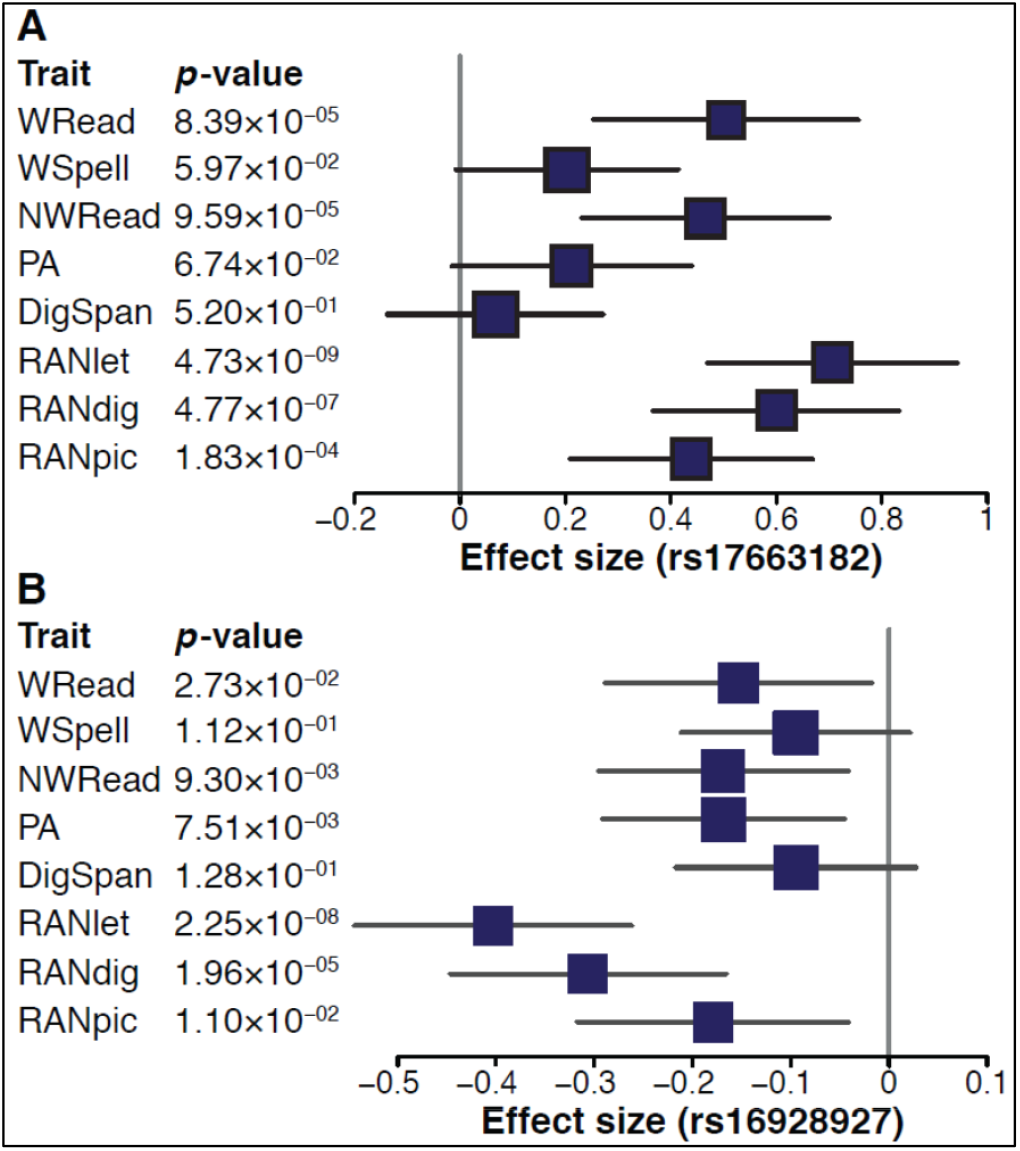
Forest plots of associations of **a)** rs17663182 (18q12.2) and **b)** rs16928927 (8q12.3) with the different traits analysed in the study. Effect sizes (β) refer to major alleles **a)** G and **b)** C, respectively.

In light of the increasing evidence implicating subcortical structures in reading and language abilities^1,35,36^, we looked up the associations of rs17663182 and rs16928927 with variability in volumes of seven different subcortical structures, which had been analysed in a previous independent GWAS^34^. After correction for multiple testing for the number of SNPs and independent traits tested (α = 7.1×10^−4^), no significant association remained (see Table S6h, i). The strongest support was observed for rs16928927 with variation in the volume of the pallidum (*p* = 6.5 × 10^−3^), where allele C was nominally associated with an increased volume (see Table S6i for details).

### Genes and SNPs previously associated with DD and related cognitive traits

12,785 variants were annotated to nine candidate genes previously implicated in dyslexia by at least two independent studies, namely *DYX1C1, DCDC2, KIAA0319, C2ORF3, MRPL19, ROBO1, GRIN2B, FOXP2*, and *CNTNAP2*. We report associations for all these variants in Table S7a-h. Among these variants, a detailed assessment of six candidate SNPs previously associated with DD or related cognitive measures in independent studies did not reveal any strong evidence of replication in our cohorts (see Table S7i).

Among variants associated with DD and related cognitive measures in previous GWAS efforts (see Table S8a-i), we identified a few nominally significant associations (*p* < 0.05) which were comparable with those reported by previous independent studies (Table S8j). The most significant associations were observed at rs10485609, with both word (A/G; MAF=12%; %; *p*-value = 2.6×10^−3^, major allele (A) β (SE) = −0.12 (0.04)) and nonword reading (*p*-value = 6.5×10^−3^, major allele (A) β (SE) = −0.1 (0.04)). These associations showed the same direction of effect as in the original report^13^.

### Gene- and pathway-based associations

Gene-level analyses of single-variant association signals in MAGMA revealed no significant associations of genes after correcting for testing of 18,033 protein-coding genes and five independent latent traits tested here (α = 5.5 ×10^−7^; see Table S9a-h). The most significant association was observed for the gene *ADCYAP1R1* (*adenylate cyclase activating polypeptide 1 receptor type I;* 7p14.3) with NWRead (Z-score = 4.6; *p* = 2×10^−6^). Similarly, also in the gene-set analysis of 1,329 canonical pathways from the MSigDB website, no pathway was significantly enriched (α = 0.01 for permutation-based enrichments, already corrected for testing of multiple pathways; see Table S10a-h). However, we found a nominally significant enrichment of associations with WSpell for genes in the BioCarta RAS pathway (Bonferroni-corrected *p* = 0.045; β(SE) = 0.64(0.16); see Table S10i for a complete list of genes leading the pathway-based association).

### Genetic overlap with neuroimaging, neurodevelopmental and neuropsychiatric traits

PRS analysis revealed the presence of a significant proportion of shared genetic variance between the different DD-related traits analysed in our GWAS and some of the neuroimaging, educational, and neuropsychiatric traits investigated in previous large GWAS studies (see Figure 5; Table S11a-c). In particular, we observed significant correlations withstanding Bonferroni correction (*p* < 8.3×10^−4^) with ADHD risk, and with educational attainment (EDUyears). The ADHD PRS was negatively associated with WRead, WSpell, NWRead, and DigSpan (at *P_T_* = 0.05: Nagelkerke’s *R*^2^ ranging from 0.004 for DigSpan to 0.007 for WRead; *p* ~ [10^−5^–10^−7^]), while EDUyears polygenic score was positively associated with WRead, WSpell, NWRead, DigSpan, and PA (at *P_T_* = 0.05: *R*^2^ ranging from 0.011 for DigSpan to 0.019 for WRead and PA; *p* ~ [10^−8^–10^−17^]). These results were confirmed at different *P_T_* thresholds (see Figure S11a-i).

**Figure 5.**
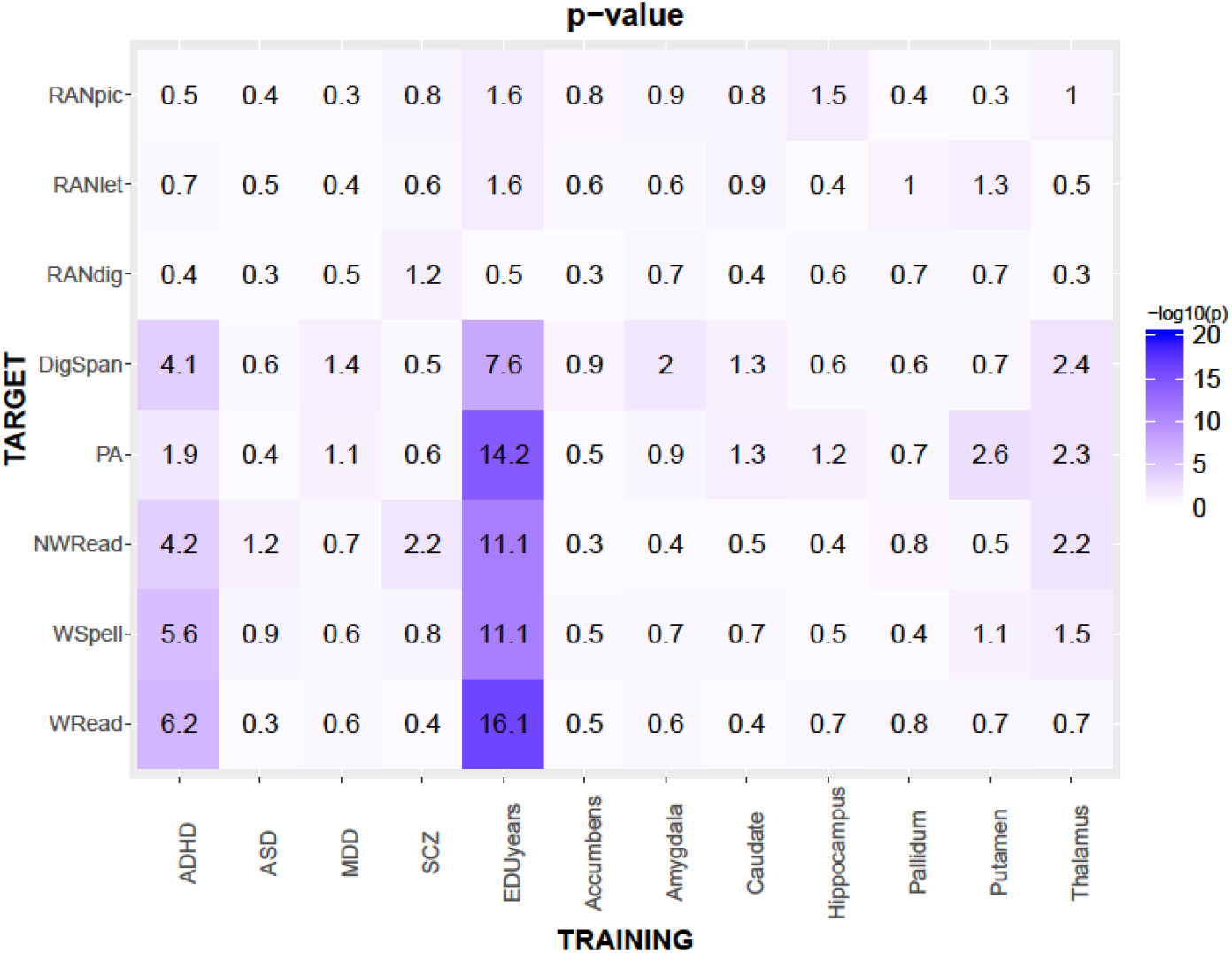
Results of the polygenic risk score (PRS) analysis on the eight traits analysed in this work (target traits), which were compared with different neuropsychiatric disorders, educational, and neuroimaging measures (training traits). In the heatmap, −log(*p*) of the *R*^2^ computed by PRSice^42^ at an association *p*-value threshold (*P_T_*) of 0.05 is reported. Complete summary statistics are reported in Tables S11a, b, c.

## Discussion

In the present study, we investigated genetic effects on eight different cognitive skills related to or underlying reading ability. We conducted a GWAS of up to 3,468 subjects from nine different countries, speaking six different languages. Hence, our study represents the richest GWAS in the field in terms of phenotypes investigated, as well as countries and languages involved.

We identified a genome-wide significant effect on rapid automatized naming of letters (RANlet). Rapid naming reflects the automaticity of visual-auditory processing necessary for a successful word decoding process and accounts for a significant proportion of variance in word reading ability, especially reading fluency, which is independent of the well-established language and phonological processes implicated in reading^51^. Moreover, RAN is considered as an excellent predictor of reading fluency and is used in kindergarten to identify children at risk of dyslexia^52^.

The most significant association signal with RANlet was observed for rs17663182, a variant located within *MIR924HG* (18q12.2; *micro-RNA 924 host gene*, or *LINC00669)*. Additional significant associations were detected in the same region for other variants, all in high LD with the lead SNP, which suggests that they identified the same genetic effect on RANlet. This observation was supported by the absence of strong independent genetic effects on RANlet within a 100 kb window surrounding the strongest signal at rs17663182. An extensive lookup of the lead variants found within this region in common online gene expression databases – including the Genotype-Tissue Expression portal (GTEx; http://www.gtexportal.org/home/)^53^, the Brain eQTL Almanac (Braineac; http://www.braineac.org/)^54^, the Blood eQTL browser (http://genenetwork.nl/bloodeqtlbrowser/)^55^, and the seeQTL database (http://www.bios.unc.edu/research/genomicsoftware/seeQTL)^56^ – revealed weak evidence of expression quantitative trait loci (eQTL) involving rs17663182 and neighboring associated SNPs. Braineac reports nominally significant eQTL effects (*p*-value < 0.05) for these SNPs on *MIR924HG* expression in the occipital cortex, thalamus, and substantia nigra. In addition, HaploReg v4.1 (http://archive.broadinstitute.org/mammals/haploreg/haploreg.php)^57^ indicated the presence of histone marks usually associated with transcriptional activity in the same region, such as H3K4me1, H3K27ac, and H3K9ac^58^. To the best of our knowledge, no regulatory role is known for *MIR924HG* and *MIR924* has not been functionally characterized so far. Nonetheless, the significant associations on 18q12.2 represent an interesting genetic effect for three main reasons:

First and foremost, evidence of genetic linkage to dyslexia-related cognitive traits has been reported for this region in previous studies, although not always reaching statistical significance^59–62^. In a genome-wide linkage analysis of a German cohort partly overlapping with our AGS dataset, a linkage peak to a principal component of RAN scores was observed in a region encompassing the microsatellite marker *D18S1102*, located ~2.1 Mb downstream of rs17663182^62^. Similarly, a linkage signal was later reported for the same marker with a composite RAN score, in a Dutch sib-pair sample. However, this association was weaker after including parents of the sib-pairs in the analysis^59^. Early evidence for linkage in 18q12 has been reported with word reading and orthographic coding, in samples partially overlapping with our Colorado and UK datasets^60,61^. In line with these findings, rs17663182 showed associations with traits other than RANlet in our analysis, including RANdig, RANpic, WRead, and NWRead (discussed below). It would be tempting to connect the linkage signals mentioned above with the SNP associations at rs17663182, but it is important to point out that this association likely represents only a small fraction of these linkage signals or even a distinct genetic effect, because linkage and association analyses tend to detect different effects^63^.

Second, a search for binding sites through the online database TargetScanHuman (http://www.targetscan.org/)^64^ allowed us to identify a series of interesting candidate target genes which *MIR924* could regulate. These include candidate susceptibility genes for dyslexia such as *MRPL19, KIAA0319L*, and *CELF4*, although these did not show the highest predicted binding scores to *MIR924* (cumulative weighted context++ scores −0.08, −0.07 and −0.04; ranked 1,615, 1,626 and 2,146 over 3,472 potential targets, respectively). Of note, the closest protein-coding gene to the associated SNPs on 18q12.2 is *CELF4* (positioned ~1.6 Mb downstream), where *D18S1102* is located. *CELF4* is highly expressed in the fetal brain and has been previously implicated in neurodevelopmental and behavioral anomalies through a haploinsufficiency mechanism^65^, although a previous candidate SNP association analysis found no major genetic effects on reading traits in this gene^61^.

Third, *MIR924HG* is expressed in a number of cancer cell lines, but consistently in samples representing iPS differentiation into neurons, according to the FANTOM5 miRNA promoter analysis^66^. This is interesting in the context that at least three dyslexia candidate genes (namely *DCDC2, DYX1C1* and *KIAA0319)* have been implicated in regulating neuronal migration and cilia functions in model systems^8^.

In the analysis of RANlet, we observed an additional association approaching genome-wide significance at rs16928927 (8q12.3). This intronic variant is located within *NKAIN3* (*Na+/K+ transporting ATPase interacting 3*), a gene which is widely and specifically expressed in the brain: in the FANTOM5 Zenbu database (http://fantom.gsc.riken.jp/zenbu/) it shows the highest expression in fetal temporal lobe, in newborn and adult hippocampal regions, and a high level of expression in all parts of the forebrain throughout development^67^. This evidence supports the importance of *NKAIN3* for neuronal function^68^ and suggests it may have a specific role in central nervous system development. Moreover, *NKAIN3* is coexpressed at the protein level with *FOXP2* – a gene previously implicated in speech, language, and reading abilities^69^ – as well as with *CELF4*, both at the transcriptional^53^ and at the translational level (the Human Integrated Protein Expression Database, available at http://www.genecards.org/)^70^. The two independent association signals on 18q12.2 and 8q12.3 might, therefore, share a common biological link with RANlet, mediated by *CELF4*. However, our SNP×SNP interaction analysis on rs17663182 and rs16928927 did not reveal any significant epistatic effect of these two variants on RANlet. The reason for this lack of support might be either the absence of an actual interaction between the two variants tested – which could still independently act in an additive manner – or that the variants are not directly causative, in which case our interaction analysis would be underpowered.

Of note, both our lead SNPs showed associations with different cognitive measures analysed in this study, especially with RAN traits. This multi-trait association trend is particularly noticeable for rs17663182, which showed convincing evidence of influence even beyond the RAN domain, extending to reading abilities. However, a formal pleiotropy test on both variants did not reveal any significant effect specific to cognitive traits other than RANlet. This suggests that these variants likely exert their genetic influence on the common phenotypic variance underlying these traits, with different magnitude of effect on each measure.

Despite the biological appeal of the top association signals mentioned above, an imaging genetic follow up of these SNPs on variation in seven different subcortical volumes previously analysed in a large independent GWAS^34^ did not reveal any significant association. Considering the sample size of the neuroimaging genetic analysis (N~ 13,000), we deem it unlikely that this lack of support is caused by a lack of power of the analysis. However, this negative result does not rule out genetic effects of the RANlet-associated variants on other brain structures involved in reading networks, such as the inferior frontal gyrus and the temporal and parietal gyri. These potential associations should be tested in the future, as was previously done for other variants associated with reading-related traits^71,72^.

Another interesting finding of our study is the significant genetic overlap that the reading traits analysed showed with educational attainment (EDUyears) and ADHD. Educational attainment was already reported to share a significant proportion of genetic variance with word reading ability^73,74^. In a PRS analysis comparing educational attainment with reading efficiency and comprehension, the same EDUYears score used in the present study^43^ accounted for 2.1% (at age 7) to 5.1% (at age 14) of the variance in such reading measures in a UK sample (N=5,825), and this association remained significant even after accounting for general cognitive ability and socioeconomic status^73^. More recently, Luciano and colleagues^74^ used the results of a previous GWAS on reading and language-related traits^11^ to test genetic correlations with several health, socioeconomic, and brain structure measures collected in adults from the UK (maximal N=111,749; age range 40-69 years). Polygenic scores increasing these traits – namely word reading, nonword repetition, and a reading-spelling score – were all positively associated with a binary index of educational attainment (college or university degree)^74^. In our paper, we replicate these findings by reporting that variants nominally associated with EDUyears explain almost 2% of the total variance in WRead and extend the evidence of genetic overlap to cognitive predictors of dyslexia risk such as PA and DigSpan. Our PRS analysis also revealed negative correlations of WRead, WSpell, NWRead, and DigSpan with an ADHD polygenic risk score^44^, suggesting the presence of a partly shared genetic basis between reading traits and ADHD risk. This long-standing hypothesis, originally supported by behavioral genetics studies of twins^75–77^, is therefore corroborated here by genome-wide genetic data.

To conclude, in the present study we report a genome-wide significant association of a variant within *MIR924HG* with RANlet, one of the best universal predictors of reading fluency across all known orthographies^78^. Our results tentatively suggest a role of this gene in the genetic etiology and neurobiology of dyslexia. Among the strengths of our study are the variety of continuous cognitive traits analysed and the relative homogeneity of phenotypic assessment and recruitment criteria of our datasets, which are fundamental to improve statistical power. Indeed, most of our samples were collected in the context of a large international consortium for studying the neurobiological/genetic basis of dyslexia (Neurodys), whose main purpose is to homogenize traits and datasets to allow for comparable analyses across different countries^19,21^. Our analyses also have some limitations, such as the absence of a follow-up cohort to replicate the genome-wide significant associations detected, as well as the relatively low sample size, compared to GWAS studies published so far in other fields^79^. The variety of the languages tested might be considered as another potential limitation of this study. Language transparency has been reported to affect the predictive power of dyslexia risk for cognitive traits such as RAN and PA, which is more pronounced in more complex orthographies^21^. Therefore, it may be hypothesized that the magnitude of genetic effects on such traits may vary depending on the transparency of the language analysed. Although the effect sizes of our most significant associations do not show any apparent relation with the transparency of the orthographies involved in the present study (see Figure 3), our analysis as presented here was designed to identify genetic effects common to and identical across language complexities, and further studies are warranted to test the specific hypothesis mentioned above. Overall, this study represents an early step of one of the largest international collaborations aimed at clarifying the genetic basis of reading abilities and disabilities, which will hopefully contribute to identify the causes of dyslexia in the future.

## Author contributions

A.G., T.F.M.A., N.M.S., D.C. and K.M. contributed to genotype QC and imputation, and to phenotype QC. A.G. carried out statistical analyses. A.G., T.F.M.A., B.M.M. and G.S.K. wrote the paper, with additional contributions by all the co-authors. All the co-authors contributed to collection, phenotypic assessment and genotyping of the datasets included in the present study. B.M.M. and G.S.K. supervised the present work.

## Acknowledgements

A.G. and T.F.M.A. were supported by the Munich Cluster for Systems Neurology (SyNergy). S.P. is a Royal Society University research Fellow. B.M.M., C.F., B.S.P. and S.E.F. are supported by the Max Planck Society. A.P.M. is a Wellcome Senior Fellow in Basic Biomedical Science (WT098017). F.R. is supported by Agence Nationale de la Recherche (ANR-06-NEURO-019-01, ANR-10-LABX-0087 IEC, ANR-10-IDEX-0001-02 PSL, ANR-11-BSV4-014-01), European Commission (LSHM-CT-2005-018696), Ville de Paris.

We would also like to acknowledge our project partners Catherine Billard, Caroline Bogliotti, Vanessa Bongiovanni, Laure Bricout, Camille Chabernaud, Yves Chaix, Isabelle Comte-Gervais, Florence Delteil-Pinton, Jean-François Démonet, Florence George, Christophe-Loïc Gérard, Stéphanie Iannuzzi, Marie Lageat, Marie-France Leheuzey, Marie-Thérèse Lenormand, Marion Liébert, Emilie Longeras, Emilie Racaud, Isabelle Soares-Boucaud, Sylviane Valdois, Nadège Villiermet, Johannes Ziegler.

## References

1 Mascheretti, S. et al. Neurogenetics of developmental dyslexia: from genes to behavior through brain neuroimaging and cognitive and sensorial mechanisms. Translational psychiatry 7, e987, doi:10.1038/tp.2016.240 (2017).

2 Raskind, W. H., Peter, B., Richards, T., Eckert, M. M. & Berninger, V. W. The Genetics of Reading Disabilities: From Phenotypes to Candidate Genes. Frontiers in Psychology 3, 601, doi:10.3389/fpsyg.2012.00601 (2012).

3 Willcutt, E. G. et al. Etiology and neuropsychology of comorbidity between RD and ADHD: The case for multiple-deficit models. Cortex 46, 1345–1361, doi:https://doi.org/10.1016/j.cortex.2010.06.009 (2010).

4 Davis, C. J. et al. Etiology of reading difficulties and rapid naming: the Colorado Twin Study of Reading Disability. Behavior genetics 31, 625–635 (2001).

5 Gayan, J. & Olson, R. K. Genetic and environmental influences on orthographic and phonological skills in children with reading disabilities. Developmental neuropsychology 20, 483–507, doi:10.1207/s15326942dn2002_3 (2001).

6 Gayan, J. & Olson, R. K. Genetic and environmental influences on individual differences in printed word recognition. Journal of experimental child psychology 84, 97–123 (2003).

7 Carrion-Castillo, A., Franke, B. & Fisher, S. E. Molecular genetics of dyslexia: an overview. Dyslexia (Chichester, England) 19, 214–240, doi:10.1002/dys.1464 (2013).

8 Kere, J. The molecular genetics and neurobiology of developmental dyslexia as model of a complex phenotype. Biochemical and biophysical research communications 452, 236–243, doi:10.1016/j.bbrc.2014.07.102 (2014).

9 Eicher, J. D. et al. Genome-wide association study of shared components of reading disability and language impairment. Genes, brain, and behavior 12, 792–801, doi:10.1111/gbb.12085 (2013).

10 Gialluisi, A. et al. Genome-wide screening for DNA variants associated with reading and language traits. Genes, brain, and behavior 13, 686–701, doi:10.1111/gbb.12158 (2014).

11 Luciano, M. et al. A genome-wide association study for reading and language abilities in two population cohorts. Genes, Brain and Behavior 12, 645–652, doi:10.1111/gbb.12053 (2013).

12 Field, L. L. et al. Dense-map genome scan for dyslexia supports loci at 4q13, 16p12, 17q22; suggests novel locus at 7q36. Genes, brain, and behavior 12, 56–69, doi:10.1111/gbb.12003 (2013).

13 Meaburn, E. L., Harlaar, N., Craig, I. W., Schalkwyk, L. C. & Plomin, R. Quantitative trait locus association scan of early reading disability and ability using pooled DNA and 100K SNP microarrays in a sample of 5760 children. Molecular psychiatry 13, 729–740, doi:10.1038/sj.mp.4002063 (2008).

14 Truong, D. et al. Multivariate genome-wide association study of rapid automatized naming and rapid alternating stimulus in Hispanic and African American youth. bioRxiv, doi:10.1101/202929 (2017).

15 Carrion-Castillo, A. et al. Association analysis of dyslexia candidate genes in a Dutch longitudinal sample. European journal of human genetics: EJHG 25, 452–460, doi:10.1038/ejhg.2016.194 (2017).

16 Willcutt, E. G., Pennington, B. F., Olson, R. K., Chhabildas, N. & Hulslander, J. Neuropsychological analyses of comorbidity between reading disability and attention deficit hyperactivity disorder: in search of the common deficit. Developmental neuropsychology 27, 35–78, doi:10.1207/s15326942dn2701_3 (2005).

17 Brandler, W. M. et al. Common variants in left/right asymmetry genes and pathways are associated with relative hand skill. PLoS genetics 9, e1003751, doi:10.1371/journal.pgen.1003751 (2013).

18 Gialluisi, A. et al. Investigating the effects of copy number variants on reading and language performance. Journal of neurodevelopmental disorders 8, 17, doi:10.1186/s11689-016-9147-8 (2016).

19 Becker, J. et al. Genetic analysis of dyslexia candidate genes in the European cross-linguistic NeuroDys cohort. European journal of human genetics: EJHG 22, 675-680, doi:10.1038/ejhg.2013.199 (2014).

20 Moll, K. et al. Cognitive mechanisms underlying reading and spelling development in five European orthographies. Learning and Instruction 29, 65–77, doi:https://doi.org/10.1016/j.learninstruc.2013.09.003 (2014).

21 Landerl, K. et al. Predictors of developmental dyslexia in European orthographies with varying complexity. Journal of child psychology and psychiatry, and allied disciplines 54, 686–694, doi:10.1111/jcpp.12029 (2013).

22 Chang, C. C. et al. Second-generation PLINK: rising to the challenge of larger and richer datasets. GigaScience 4, 7, doi:10.1186/s13742-015-0047-8 (2015).

23 Andlauer, T. F. et al. Novel multiple sclerosis susceptibility loci implicated in epigenetic regulation. Science advances 2, e1501678, doi: 10.1126/sciadv.1501678 (2016).

24 The Genomes Project, C. A global reference for human genetic variation. Nature 526, 68, doi:10.1038/nature15393 (2015).

25 Delaneau, O., Zagury, J.-F. & Marchini, J. Improved whole-chromosome phasing for disease and population genetic studies. Nat Meth 10, 5–6, doi:10.1038/nmeth.2307 (2013).

26 Howie, B. N., Donnelly, P. & Marchini, J. A flexible and accurate genotype imputation method for the next generation of genome-wide association studies. PLoS genetics 5, e1000529, doi:10.1371/journal.pgen.1000529 (2009).

27 Lippert, C. et al. FaST linear mixed models for genome-wide association studies. Nat Meth 8, 833–835, doi: 10.1038/nmeth.1681 (2011).

28 Han, B. & Eskin, E. Random-Effects Model Aimed at Discovering Associations in Meta-Analysis of Genome-wide Association Studies. The American Journal of Human Genetics 88, 586–598, doi:10.1016/j.ajhg.2011.04.014.

29 Li, J. & Ji, L. Adjusting multiple testing in multilocus analyses using the eigenvalues of a correlation matrix. Heredity 95, 221–227, doi:10.1038/sj.hdy.6800717 (2005).

30 Aulchenko, Y. S., Ripke, S., Isaacs, A. & van Duijn, C. M. GenABEL: an R library for genome-wide association analysis. Bioinformatics (Oxford, England) 23, 1294-1296, doi:10.1093/bioinformatics/btm108 (2007).

31 (2005).

32 Langfelder, P. & Horvath, S. WGCNA: an R package for weighted correlation network analysis. BMC bioinformatics 9, 559, doi:10.1186/1471-2105-9-559 (2008).

33 Willer, C. J., Li, Y. & Abecasis, G. R. METAL: fast and efficient meta-analysis of genomewide association scans. Bioinformatics (Oxford, England) 26, 2190–2191, doi:10.1093/bioinformatics/btq340 (2010).

34 Hibar, D. P. et al. Common genetic variants influence human subcortical brain structures. Nature 520, 224–229, doi:10.1038/nature14101 (2015).

35 Eicher, J. D. & Gruen, J. R. Imaging-genetics in dyslexia: Connecting risk genetic variants to brain neuroimaging and ultimately to reading impairments. Molecular Genetics and Metabolism 110, 201–212, doi:https://doi.org/10.1016/j.ymgme.2013.07.001 (2013).

36 Krishnan, S., Watkins, K. E. & Bishop, D. V. Neurobiological Basis of Language Learning Difficulties. Trends in cognitive sciences 20, 701–714, doi:10.1016/j.tics.2016.06.012 (2016).

37 Bates, T. C. et al. Genetic variance in a component of the language acquisition device: ROBO1 polymorphisms associated with phonological buffer deficits. Behavior genetics 41, 50–57, doi:10.1007/s10519-010-9402-9 (2011).

38 Tran, C. et al. Association of the ROBO1 gene with reading disabilities in a family-based analysis. Genes, brain, and behavior 13, 430–438, doi: 10.1111/gbb. 12126 (2014).

39 Peter, B. et al. Replication of CNTNAP2 association with nonword repetition and support for FOXP2 association with timed reading and motor activities in a dyslexia family sample. Journal of neurodevelopmental disorders 3, 39–49, doi:10.1007/s11689-010-9065-0 (2011).

40 Mascheretti, S. et al. GRIN2B mediates susceptibility to intelligence quotient and cognitive impairments in developmental dyslexia. Psychiatric genetics 25, 9–20, doi:10.1097/ypg. 0000000000000068 (2015).

41 de Leeuw, C. A., Mooij, J. M., Heskes, T. & Posthuma, D. MAGMA: generalized gene-set analysis of GWAS data. PLoS computational biology 11, e1004219, doi:10.1371/journal.pcbi.1004219 (2015).

42 Euesden, J., Lewis, C. M. & O’Reilly, P. F. PRSice: Polygenic Risk Score software. Bioinformatics (Oxford, England) 31, 1466–1468, doi:10.1093/bioinformatics/btu848 (2015).

43 Okbay, A. et al. Genome-wide association study identifies 74 loci associated with educational attainment. Nature 533, 539–542, doi:10.1038/nature17671 (2016).

44 Demontis, D. et al. Discovery Of The First Genome-Wide Significant Risk Loci For ADHD. bioRxiv, doi: 10.1101/145581 (2017).

45 Meta-analysis of GWAS of over 16,000 individuals with autism spectrum disorder highlights a novel locus at 10q24.32 and a significant overlap with schizophrenia. Molecular Autism 8, 21, doi:10.1186/s13229-017-0137-9 (2017).

46 Ripke, S. et al. A mega-analysis of genome-wide association studies for major depressive disorder. Molecular psychiatry 18, 497–511, doi:10.1038/mp.2012.21 (2013).

47 Biological insights from 108 schizophrenia-associated genetic loci. Nature 511, 421–427, doi:10.1038/nature 13595 (2014).

48 Russell, G. & Pavelka, Z. in Recent Advances in Autism Spectrum Disorders - Volume I (ed Michael Fitzgerald) Ch. 17 (InTech, 2013).

49 Mugnaini, D., Lassi, S., La Malfa, G. & Albertini, G. Internalizing correlates of dyslexia. World Journal of Pediatrics 5, 255–264, doi:10.1007/s12519-009-0049-7 (2009).

50 Whitford, V., O’Driscoll, G. A. & Titone, D. Reading deficits in schizophrenia and their relationship to developmental dyslexia: A review. Schizophrenia Research, doi:https://doi.org/10.1016/j.schres.2017.06.049 (2017).

51 Wolf, M. B., Patricia Greig. The double-deficit hypothesis for the developmental dyslexias. Journal of Educational Psychology 91, 415–438 (1999).

52 Schatschneider, C., Fletcher, J. M., Francis, D. J., Carlson, C. D. & Foorman, B. R. Kindergarten Prediction of Reading Skills: A Longitudinal Comparative Analysis. Journal of Educational Psychology 96, 265–282, doi:10.1037/0022-0663.96.2.265 (2004).

53 The, G. C. The Genotype-Tissue Expression (GTEx) project. Nature genetics 45, 580–585, doi:10.1038/ng.2653 (2013).

54 Ramasamy, A. et al. Genetic variability in the regulation of gene expression in ten regions of the human brain. Nature neuroscience 17, 1418–1428, doi:10.1038/nn.3801 (2014).

55 Westra, H. J. et al. Systematic identification of trans eQTLs as putative drivers of known disease associations. Nature genetics 45, 1238–1243, doi:10.1038/ng.2756 (2013).

56 Xia, K. et al. seeQTL: a searchable database for human eQTLs. Bioinformatics (Oxford, England) 28, 451–452, doi:10.1093/bioinformatics/btr678 (2012).

57 Ward, L. D. & Kellis, M. HaploReg v4: systematic mining of putative causal variants, cell types, regulators and target genes for human complex traits and disease. Nucleic acids research 44, D877–881, doi:10.1093/nar/gkv1340 (2016).

58 Pérez-Lluch, S., Guigó, R. & Corominas, M. Active transcription without histone modifications. Oncotarget 6, 41401–41401 (2015).

59 de Kovel, C. G. et al. Confirmation of dyslexia susceptibility loci on chromosomes 1p and 2p, but not 6p in a Dutch sib-pair collection. American journal of medical genetics. Part B, Neuropsychiatric genetics: the official publication of the International Society of Psychiatric Genetics 147, 294–300, doi:10.1002/ajmg.b.30598 (2008).

60 Fisher, S. E. et al. Independent genome-wide scans identify a chromosome 18 quantitative-trait locus influencing dyslexia. Nature genetics 30, 86–91, doi:10.1038/ng792 (2002).

61 Scerri, T. S. et al. Identification of candidate genes for dyslexia susceptibility on chromosome 18. PloS one 5, e13712, doi:10.1371/journal.pone.0013712 (2010).

62 Schulte-Korne, G. et al. Interrelationship and familiality of dyslexia related quantitative measures. Annals of human genetics 71, 160–175, doi: 10.1111/j. 1469-1809.2006.00312.x (2007).

63 Ott, J., Kamatani, Y. & Lathrop, M. Family-based designs for genome-wide association studies. Nature Reviews Genetics 12, 465, doi:10.1038/nrg2989 (2011).

64 Agarwal, V., Bell, G. W., Nam, J.-W. & Bartel, D. P. Predicting effective microRNA target sites in mammalian mRNAs. eLife 4, e05005, doi:10.7554/eLife.05005 (2015).

65 Halgren, C. et al. Haploinsufficiency of CELF4 at 18q12.2 is associated with developmental and behavioral disorders, seizures, eye manifestations, and obesity. European journal of human genetics: EJHG 20, 1315–1319, doi:10.1038/ejhg.2012.92 (2012).

66 de Rie, D. et al. An integrated expression atlas of miRNAs and their promoters in human and mouse. Nature Biotechnology 35, 872, doi:10.1038/nbt.3947 (2017).

67 The, F. C., the, R. P. & Clst. A promoter-level mammalian expression atlas. Nature 507, 462, doi:10.1038/nature13182 (2014).

68 Gorokhova, S., Bibert, S., Geering, K. & Heintz, N. A novel family of transmembrane proteins interacting with β subunits of the Na,K-ATPase. Human Molecular Genetics 16, 2394–2410, doi:10.1093/hmg/ddm167 (2007).

69 Graham, S. A. & Fisher, S. E. Understanding Language from a Genomic Perspective. Annual review of genetics 49, 131–160, doi:10.1146/annurev-genet-120213-092236 (2015).

70 Fishilevich, S. et al. Genic insights from integrated human proteomics in GeneCards. Database: the journal of biological databases and curation 2016, doi:10.1093/database/baw030 (2016).

71 Eicher, J. D. et al. Dyslexia and language impairment associated genetic markers influence cortical thickness and white matter in typically developing children. Brain Imaging and Behavior 10, 272–282, doi:10.1007/s11682-015-9392-6 (2016).

72 Gialluisi, A., Guadalupe, T., Francks, C. & Fisher, S. E. Neuroimaging genetic analyses of novel candidate genes associated with reading and language. Brain and language 172, 9–15, doi:10.1016/j.bandl.2016.07.002 (2017).

73 Selzam, S. et al. Genome-Wide Polygenic Scores Predict Reading Performance Throughout the School Years. Scientific Studies of Reading 21, 334–349, doi:10.1080/10888438.2017.1299152 (2017).

74 Luciano, M. et al. Single Nucleotide Polymorphisms Associated with Reading Ability Show Connection to Socio-Economic Outcomes. Behavior genetics, doi:10.1007/s10519-017-9859-x (2017).

75 Greven, C. U., Harlaar, N., Dale, P. S. & Plomin, R. Genetic Overlap between ADHD Symptoms and Reading is largely Driven by Inattentiveness rather than Hyperactivity-Impulsivity. Journal of the Canadian Academy of Child and Adolescent Psychiatry 20, 6–14 (2011).

76 Willcutt, E. G., Pennington, B. F. & DeFries, J. C. Twin study of the etiology of comorbidity between reading disability and attention-deficit/hyperactivity disorder. American journal of medical genetics 96, 293–301 (2000).

77 Willcutt, E. G., Pennington, B. F., Olson, R. K. & DeFries, J. C. Understanding comorbidity: A twin study of reading disability and attention-deficit/hyperactivity disorder. American Journal of Medical Genetics Part B: Neuropsychiatric Genetics 144B, 709–714, doi:10.1002/ajmg.b.30310 (2007).

78 Norton, E. S. & Wolf, M. Rapid automatized naming (RAN) and reading fluency: implications for understanding and treatment of reading disabilities. Annual review of psychology 63, 427–452, doi:10.1146/annurev-psych-120710-100431 (2012).

79 Locke, A. E. et al. Genetic studies of body mass index yield new insights for obesity biology. Nature 518, 197, doi:10.1038/nature14177 (2015).

80 Wechsler, D. The Wechsler intelligence scale for children. 3rd ed. edn, (London: The Psychological Corporation., 1992).

81 Wechsler, D. Wechsler intelligence scale for children. 4th ed. edn, (San Antonio, TX: Psychological Corporation., 2003).

82 Wechsler, D. Manual for the Wechsler Intelligence Scale for Children – Revised., (The Psychological Corporation, New York, NY., 1974).

83 Wechsler, D. Manual for the Wechsler Adult Intelligence Scale – Revised., (Psychological Corporation, New York, NY., 1981).

84 Elliot, C. D., Murray, D.J. & Pearson, L.S. The British Ability Scales., (NFER, Slough, UK., 1979).

